# No-known-vector flaviviruses exhibit diverse replication and virulence phenotypes in mice

**DOI:** 10.64898/2026.06.11.731570

**Authors:** Mikhala E. Dorminey, Joshua R. Nielsen, Wes Sanders, Nathaniel J. Moorman, Helen M. Lazear

## Abstract

The *Orthoflavivirus* genus (flaviviruses) includes globally significant arboviruses, which cycle between arthropod vectors (mosquitoes and ticks) and vertebrate hosts. In contrast, no-known-vector flaviviruses (NKVFVs) have been isolated from rodents and bats, but not arthropods, so are thought to spread by vector-independent routes. However, little is known about the host range and pathogenic mechanisms of these viruses. To evaluate NKVFV pathogenesis, we infected wild-type, *Ifnar1^−/−^*, and *Ifnar1^−/−^ Ifngr1^−/−^* mice by footpad inoculation with 12 NKVFVs: Entebbe bat virus (ENTV), Sokuluk virus (SOKV), Yokose virus (YOKV), Modoc virus (MODV), Apoi virus (APOIV), Jutiapa virus (JUTV), Sal Vieja virus (SVV), Dakar bat virus (DBV), Rio Bravo virus (RBV), Montana myotis leukoencephalitis virus (MMLV), Phnom Penh bat virus (PPBV), and Tamana bat virus (TABV). We compared these NKVFVs to the mosquito-borne Zika virus and Kedougou virus and to the tick-borne Langat virus and Kadam virus. We monitored disease signs and measured viremia. 7 NKVFVs (ENTV, SOKV, YOKV, MODV, APOIV, DBV, RBV) were virulent in *Ifnar1*^−/−^ mice, causing 100% lethality within 9 dpi, accompanied by viremia. All viruses tested were virulent in *Ifnar1^−/−^ Ifngr1^−/−^* mice and produced greater viremia compared to *Ifnar1*^−/−^ mice. No viremia or disease signs were detected in wild-type mice. We further evaluated RBV, MMLV, and PPBV replication in mouse primary fibroblasts and bone marrow-derived macrophages, as well as in cell lines from three bat species. Altogether, our results provide new information about the virulence and replication phenotypes of NKVFVs, supporting future studies investigating NKVFV-specific and pan-flavivirus pathogenic mechanisms.

**IMPORTANCE:** Flaviviruses that cause human disease are transmitted by mosquitoes and ticks (e.g. West Nile virus, yellow fever virus, tick-borne encephalitis virus). But there are related flaviviruses that are not known to infect arthropods, so are thought to spread by vector-independent routes (no-known-vector flaviviruses, NKVFVs). Not much is known about NKVFVs, but they provide an opportunity to understand flavivirus replication, tropism, and pathogenesis more broadly. We evaluated a panel of 12 NKVFVs for their ability to cause disease in mice with and without antiviral interferon responses as well as their replication in mouse cells. Our findings provide new information about these under-studied viruses and demonstrate which mouse models may be appropriate to use for further studies with NKVFVs.

## INTRODUCTION

The *Orthoflavivirus* genus (flaviviruses) includes globally significant arboviruses such as West Nile virus, dengue virus, yellow fever virus, and tick-borne encephalitis virus, and the need for these viruses to cycle between arthropod vectors and vertebrate hosts underlies their ecology, epidemiology, and evolution (1). Flaviviruses that are known human pathogens are transmitted by mosquitoes or ticks (mosquito-borne flaviviruses, MBFV; and tick-borne flaviviruses, TBFV). *Orthoflavivirus* also includes viruses that are vertically transmitted in mosquitoes, but do not infect vertebrates (insect-specific flaviviruses, ISFV), as well as viruses that are found in vertebrates but have no known arthropod vector (no-known vector flaviviruses, NKVFV) (2–6). In general, flavivirus transmission mechanisms are reflected in the phylogeny of these viruses, with MBFV, TBFV, ISFV, and NKVFV forming several distinct genetic groupings, though vector and host tropism are not monophyletic (7). For example, NKVFVs are found in three distinct clades: one within the MBFV group, one adjacent to TBFVs, and one forming a separate genus (4–6). The relationships among these genetic groupings and exceptions to these trends suggest that over the course of flavivirus evolution, there have been multiple instances of flaviviruses expanding or restricting their host range and transmission mechanisms, creating the potential for the emergence of new human or animal pathogens (8). Understanding the mechanisms that govern flavivirus vector and host tropism and how that determines transmission mechanisms is important for evaluating the risk of future emerging viruses. Furthermore, understanding the tissue and species tropism, and underlying virus-host interactions, of flaviviruses with diverse host ranges and transmission mechanisms can provide insight into the replication and pathogenesis of flaviviruses that already are pathogens of humans, livestock, and wildlife.

NKVFVs have been isolated from rodents and bats around the world, but little is known about their ecology, cell biology, and pathogenesis (2). NKVFV pathogenesis has been evaluated in various models, including mice, hamsters, non-human primates, chicken embryos, rabbits, and bats (9–14). However, prior studies have assessed limited sets of NKVFVs and have not generally employed modern transgenic mouse lines, which support mechanistic studies of the host mechanisms that control viral tropism and pathogenesis. We sought to evaluate a large panel of NKVFVs (12 of the 17 viruses reported) using common mouse models, with the goal of defining the pathogenic features of these viruses and identifying mouse models that could support research in the context of a future emergence or outbreak (4–6). We used wild-type C57BL/6 mice and mice lacking type I interferon (IFN-αβ) signaling (Ifnar*1^−/−^*) or type I and type II (IFN-γ) signaling (*Ifnar1^−/−^Ifngr1^−/−^*) as well as primary cells derived from these mice. We further evaluated replication of bat-origin NKVFVs in bat cell lines, to better model the interaction of these viruses with their bat hosts. Altogether, this work provides the first comprehensive comparison of diverse NKVFVs in mouse models, expanding our understanding of these viruses and providing systems for investigating the pathogenic mechanisms of flaviviruses broadly.

## RESULTS

### Complete genome sequences and phylogenetic comparison of orthoflaviviruses

To confirm sequences of laboratory viral stocks, we deep sequenced stocks of Aedes flavivirus, Apoi virus, Aroa virus, Banzi virus, Culex flavivirus, Dakar bat virus, Iguape virus, Jutiapa virus, Kedougou virus, Modoc virus, Phnom Penh bat virus, Rio Bravo virus, Sal Vieja virus, Sokuluk virus, Tamana bat virus, Usutu virus, and Yokose virus, generating full coding sequences (**Table 1**). While full coding sequences previously were available for most of these viruses, the previously available sequence for Sal Vieja virus comprised only a partial NS5 coding sequence. We also generated a complete M and E coding sequence for Dakar bat virus strain AD D249, for which no sequence previously was available. We constructed a phylogenetic tree of orthoflaviviruses based on full coding sequences; the topology of the tree, including the separation of NKVFVs into 3 distinct clades, is consistent with prior work (**Figure 1**) (4–6).

**Figure 1.**
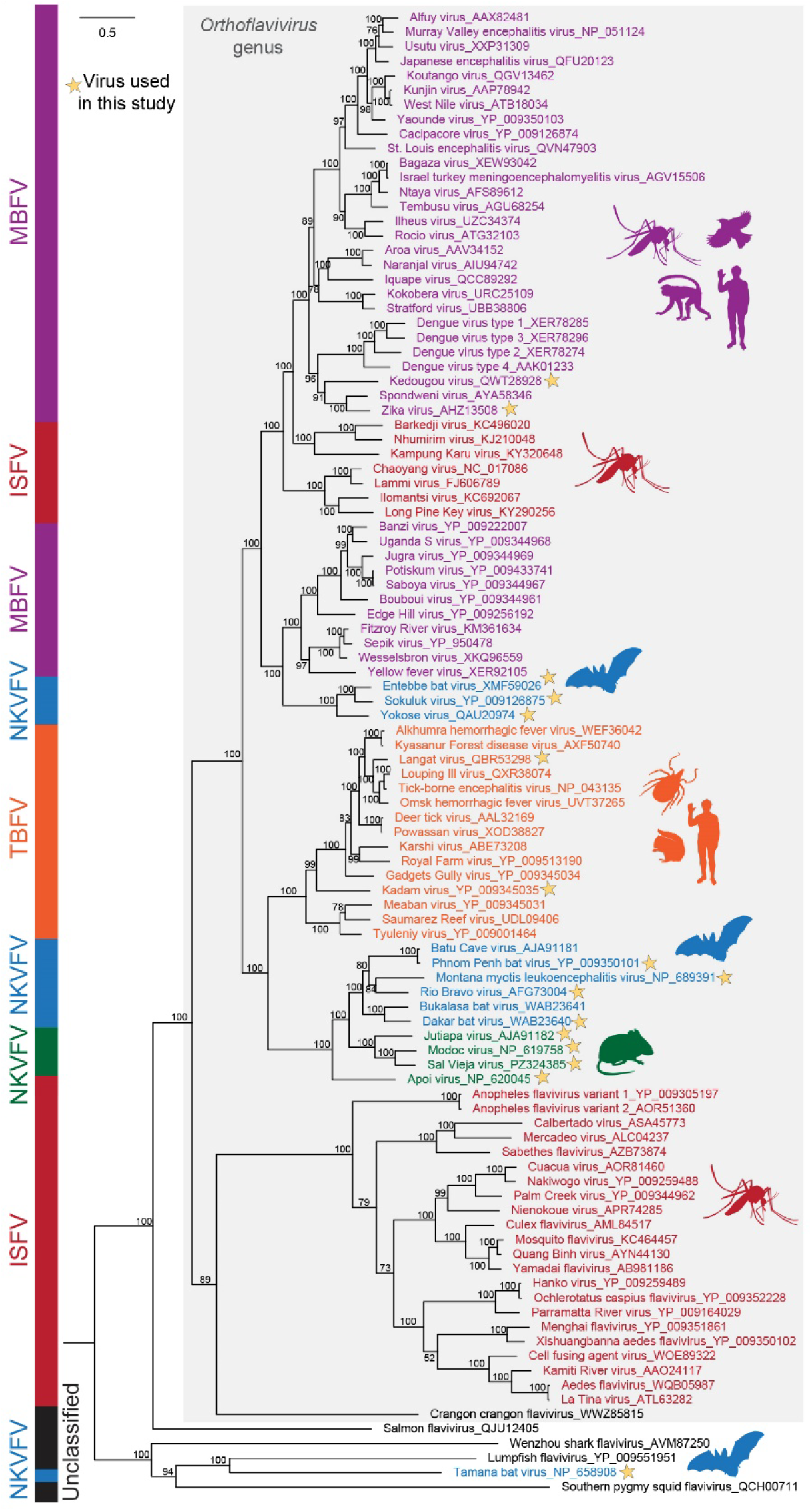
Orthoflavivirus phylogeny. Full polyprotein sequences of 102 flaviviruses were aligned using MUSCLE and a mid-rooted maximum likelihood phylogenetic tree generated; bootstrap values are indicated. Viruses are color-coded based on their vector usage: mosquito-borne (purple), tick-borne (orange), insect-specific (red), no-known-vector, bat-origin (blue), no-known-vector, rodent-origin (green), or unclassified (black). Gray shading indicates the Orthoflavivirus genus within the Flaviviridae family. Genbank accession numbers for each virus are indicated. Viruses included in this study are indicated with a star. Scale bar represents 0.5 substitutions per site. Silhouette images are from PhyloPic.org.

**Table 1:** Virus stocks sequenced in this study.

| Virus | Genbank Accession # |
| --- | --- |
| Aedes flavivirus | PZ303603 |
| Apoi virus | PZ305740 |
| Aroa virus | PZ305741 |
| Banzi virus | PZ305742 |
| Culex flavivirus | PZ305743 |
| Dakar bat virus strain BP180 | PZ305744 |
| Dakar bat virus strain AD D249 | PZ500116 |
| Iguape virus | PZ324379 |
| Jutiapa virus | PZ324380 |
| Kedougou virus | PZ324381 |
| Modoc virus | PZ324382 |
| Phnom Penh bat virus | PZ324383 |
| Rio Bravo virus | PZ324384 |
| Sal Vieja virus | PZ324385 |
| Sokuluk virus | PZ324386 |
| Tamana bat virus | PZ324387 |
| Usutu virus | PZ324389 |
| Yokose virus | PZ324388 |

### No-known-vector flavivirus infection causes variable lethality and viremia in mice

To evaluate NKVFV virulence in mice, we infected five to six week old wild-type C57BL/6, *Ifnar1^−/−^*, and *Ifnar1^−/−^Ifngr1^−/−^* mice with 1,000 focus-forming units (FFU) by subcutaneous inoculation in the footpad of 12 NKVFVs (Entebbe bat virus (ENTV), Sokuluk virus (SOKV), Yokose virus (YOKV), Apoi virus (APOIV), Modoc virus (MODV), Jutiapa virus (JUTV), Sal Vieja virus (SVV), Dakar bat virus (DBV), Montana myotis leukoencephalitis virus (MMLV), Rio Bravo virus (RBV), Phnom Penh bat virus (PPBV), and Tamana bat virus (TABV)) **(Table 2)** and evaluated lethality for 21 days (**Figure 2**). For comparison, we also infected mice with the mosquito-borne Zika virus (ZIKV) and Kedougou virus (KEDV) and the tick-borne Kadam virus (KADV) and Langat virus (LGTV) (**Figure 2**). ZIKV and LGTV caused 100% lethality in *Ifnar1^−/−^*, and *Ifnar1^−/−^Ifngr1^−/−^* mice and no lethality in wild-type mice, consistent with previous studies (15, 16), while KEDV and KADV exhibited similar virulence. None of the tested NKFVs caused lethality in wild-type mice, which is especially notable for NKVFVs that were originally isolated from rodents (e.g. APOIV, MODV). We observed varying degrees of lethality in both the *Ifnar1^−/−^* and *Ifnar1^−/−^ Ifngr1^−/−^* mice. ENTV, SOKV, YOKV, APOIV, MODV, DBV, and RBV all caused 100% lethality in *Ifnar1^−/−^* mice, whereas SVV, PPBV, and TABV each caused <25% lethality. Two additional strains of DBV (AD D249 and IBAN 8646) exhibited equivalent virulence as DBV strain BP 180 (data not shown). JUTV and MMLV caused no lethality in *Ifnar1^−/−^*mice. All viruses tested in *Ifnar1^−/−^Ifngr1^−/−^* mice (ZIKV, KADV, LGTV, JUTV, SVV, RBV, MMLV, PPBV, and TABV) caused >95% lethality. Some viruses that caused 100% lethality in *Ifnar1^−/−^* mice were not evaluated in *Ifnar1^−/−^Ifngr1^−/−^* mice (KEDV, ENTV, SOKV, YOKV, APOIV, MODV, DBV). These results demonstrate IFN-αβ signaling as a key host response restricting NKVFV virulence in mice, with IFN-γ signaling contributing additional control, consistent with findings with many vector-borne flaviviruses, including ZIKV (17, 18), Spondweni virus (19), Usutu virus (20), and Langat virus (21).

**Figure 2.**
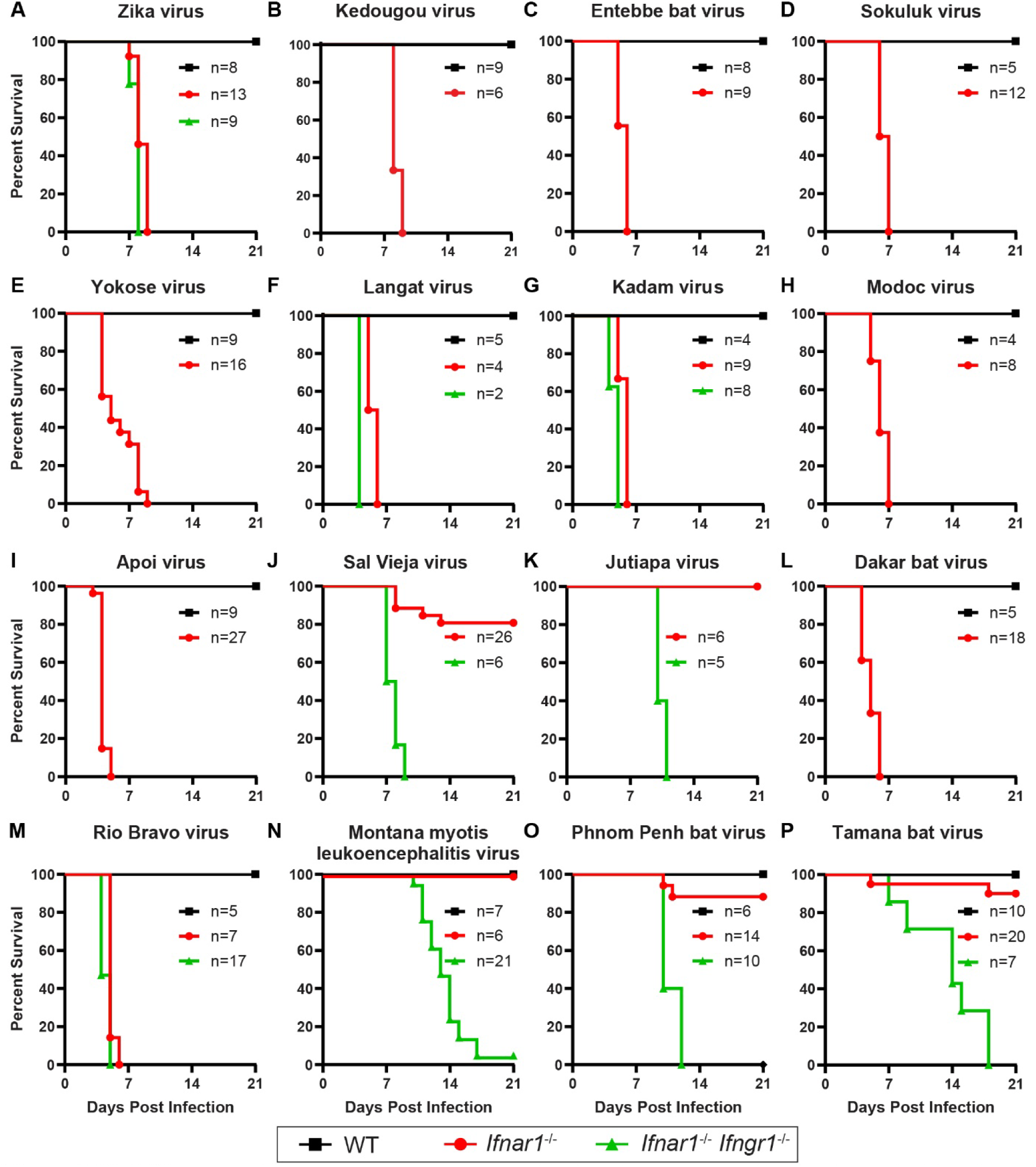
Variable survival outcomes occur after NKVFV infection in mice. Five to six-week-old male and female wild-type (WT), *Ifnar1*^−/−^, and *Ifnar1*^−/−^ *Ifngr1*^−/−^ mice were infected with 1000 focus-forming units (FFU) of the indicated virus and strain by subcutaneous inoculation in the footpad. **A.** Zika virus (ZIKV) (H/PF/2013). **B.** Kedougou virus (KEDV) (DAK AR D 14701). **C.** Entebbe bat virus (ENTV) (IL 30). **D.** Sokuluk virus (SOKV) (LEIV 400K). **E.** Yokose virus (YOKV) (Oita 36). **F.** Langat virus (LGTV) (TP-21) **G.** Kadam virus (KADV) (T 100). **H.** Modoc virus (MODV) (M 544). **I.** Apoi virus (APOIV) (Apoi). **J.** Sal Vieja virus (SVV) (78 TWM 106). **K.** Jutiapa virus (JUTV) (JG 128). **L.** Dakar bat virus (DBV) (BP 180). **M.** Rio Bravo virus (RBV) (M64). **N.** Montana myotis leukoencephalitis virus (MMLV) (B 334 A-924 BF). **O.** Phnom Penh bat virus (PPBV) (A38-69). **P.** Tamana bat virus (TABV) (TR 127154). Mice were monitored for lethality for 21 days. Data are combined from 2 to 3 experiments per virus, with the number of mice per group indicated.

**Table 2:** No-known-vector flaviviruses used in this study.

| Virus | Abbreviation (strain) | Isolation |  |  |  | References |
| --- | --- | --- | --- | --- | --- | --- |
|  |  | Year | Location | Species | Tissue |  |
| Entebbe bat virus | ENTV (IL 30) | 1957 | Uganda | *Tadarida pumila* (little free-tailed bat) | Salivary gland | (22–25) |
| Sokuluk virus | SOKV (LEIV 400K) | 1970 | USSR (Kyrgyzstan) | *Pipistrellus pipistrellus* (common pipistrelle) | Pooled samples of brain, liver, spleen, and kidney | (26) |
| Yokose virus | YOKV (Oita 36) | 1971 | Japan (Kyushu Island) | *Miniopterus fuliginosus* (eastern bent-wing bat) | Blood | (27, 28) |
| Modoc virus | MODV (M544) | 1958 | USA (California) | *Peromyscus maniculatus* (eastern deer mouse) | Lactating mammary glands | (29, 30) |
| Apoi virus | APOIV (Apoi) | 1954 | Japan (Hokkaido) | *Apodemus speciosus* (large Japanese field mouse), *Clethrionomys rufocanus* (grey red-backed vole), *Apodemus argenteus* (small Japanese field mouse) | Pooled spleens | (31) |
| Sal Vieja virus | SVV (78 TWM 106) | 1978 | USA (Texas) | *Peromyscus leucopus* (white-footed mouse) | (not specified) | (32) |
| Jutiapa virus | (JG 128) | 1968 | Guatemala (Jutiapa) | *Sigmodon hispidus* (hispid cotton rat) | Serum/plasma | (29) |
| Dakar bat virus | DBV (BP 180) | 1962 | Senegal | *Tadarida pumila* (little free-tailed bat) | Salivary gland | (33) |
| Rio Bravo virus | RBV (M64) | 1954 | USA (California) | *Tadarida brasiliensis* (Mexican free-tailed bat) | Salivary gland | (34) |
| Montana myotis leukoencephalitis virus | MMLV (B 334 A-924BF) | 1964 | USA (Montana) | *Myotis lucifugus* (little brown bat) | Salivary gland | (35) |
| Phnom Penh bat virus | PPBV (A38-69) | 1969 | Cambodia | *Cynopterus brachyotis* (lesser short-nosed fruit bat) | Salivary gland and brown fat | (36) |
| Tamana bat virus | TABV (TR 127154) | 1973 | Trinidad | *Pteronotus parnellii* (Parnell's mustached bat) | Salivary glands, saliva, and spleen | (37–39) |

To determine whether NKVFV could replicate in mice and produce viremia, we collected serial serum samples via submandibular bleed at 2, 4, and 6 days post-infection (dpi) from the same mice used in lethality studies and measured infectious viral titers by focus-forming assay (FFA) (**Figure 3**). We did not detect infectious virus in the serum of wild-type mice infected with any of the viruses tested. In *Ifnar1^−/−^* mice, we observed four distinct trends for viremia measurements over time: i) viremia peaked 2 dpi then declined (ZIKV, ENTV, SOKV, YOKV, MODV, SVV, and TABV), ii) viremia was constant across all timepoints collected (JUTV, KADV, and RBV), iii) viremia increased across all timepoints collected (APOIV, DBV, and LGTV), or iv) no viremia detected (MMLV and PPBV). Two additional strains of DBV (AD D249 and IBAN 8646) exhibited equivalent viremia as DBV strain BP 180 (data not shown). In general, the magnitude and duration of viremia was greater in *Ifnar1^−/−^ Ifngr1^−/−^* mice compared to *Ifnar1^−/−^*, with the same four trends evident: SVV and KADV produced viremia that peaked 2dpi and began to clear; ZIKV and TABV produced viremia that was constant across all timepoints collected; JUTV and RBV produced increasing viremia; and no viremia was detected from MMLV. In both *Ifnar1^−/−^*mice and *Ifnar1^−/−^ Ifngr1^−/−^* mice, some viruses produced rapid lethality that precluded an assessment of viremia trends over the full 6-day period. Overall, the relationship between viremia (**Figure 3**) and virulence (**Figure 2**) was inconsistent. For example, *Ifnar1^−/−^* mice succumbed rapidly to APOIV (mice began to succumb at 3 dpi, with 100% mortality by 5 dpi), but viremia in mice infected with APOIV at 2 dpi was no higher than some viruses that resulted in slower lethality (YOKV, TABV). Conversely, MODV produced 100% lethality by 6 to 8 dpi in *Ifnar1^−/−^* mice with similar kinetics to other viruses (e.g. ENTV, SOKV, YOKV, KADV, LGTV, DBV, RBV), but MODV viremia at 2 and 4 dpi was lower than other viruses. No viremia was detected in *Ifnar1^−/−^ Ifngr1^−/−^* mice infected with MMLV or PPBV even though infection resulted in 95-100% lethality. To increase the sensitivity of our viremia measurements, we analyzed serum from RBV, MMLV, and PPBV infected *Ifnar1^−/−^* and *Ifnar1^−/−^ Ifngr1^−/−^* by RT-qPCR. With this approach, we detected viremia at all time points (**Figure 4**). Overall, these results show that NKVFV can produce viremia in susceptible mice, but viremia does not directly correspond to virulence.

**Figure 3.**
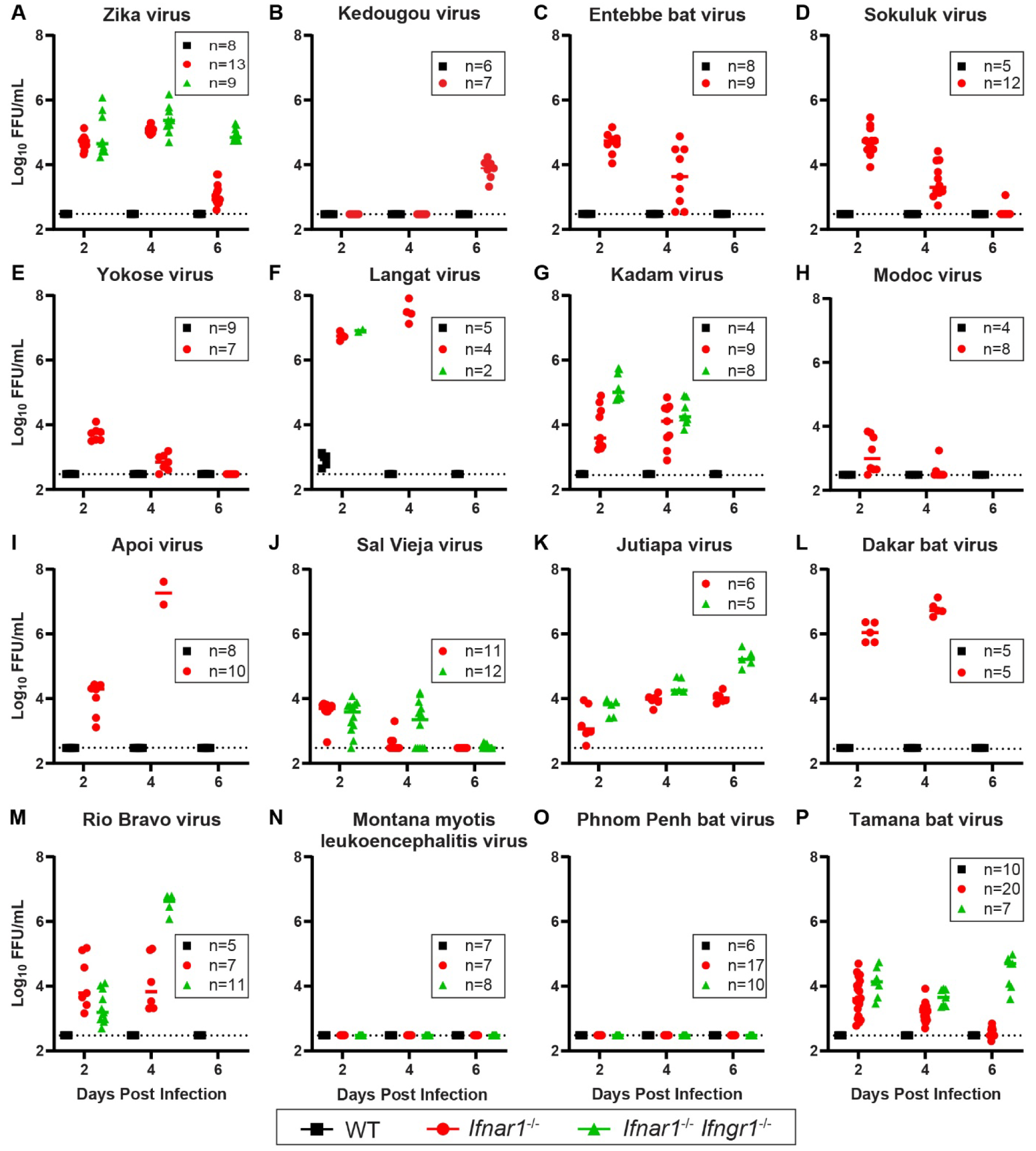
Diverse viremia profiles are produced by NKVFV infection in mice. Five to six-week-old male and female wild-type (WT), *Ifnar1*^−/−^, and *Ifnar1*^−/−^ *Ifngr1*^−/−^ mice were infected with 1000 FFU of the indicated flavivirus by subcutaneous inoculation in the footpad. **A.** Zika virus (ZIKV) (H/PF/2013). **B.** Kedougou virus (KEDV) (DAK AR D 14701). **C.** Entebbe bat virus (ENTV) (IL 30). **D.** Sokuluk virus (SOKV) (LEIV 400K). **E.** Yokose virus (YOKV) (Oita 36). **F.** Langat virus (LGTV) (TP-21) **G.** Kadam virus (KADV) (T 100). **H.** Modoc virus (MODV) (M 544). **I.** Apoi virus (APOIV) (Apoi). **J.** Sal Vieja virus (SVV) (78 TWM 106). **K.** Jutiapa virus (JUTV) (JG 128). **L.** Dakar bat virus (DBV) (BP 180). **M.** Rio Bravo virus (RBV) (M64). **N.** Montana myotis leukoencephalitis virus (MMLV) (B 334 A-924 BF). **O.** Phnom Penh bat virus (PPBV) (A38-69). **P.** Tamana bat virus (TABV) (TR 127154). Mice were bled at 2-, 4-, and 6-days post-infection and viremia was measured by focus-forming assay. Data are combined from 2 to 3 experiments per virus, with the number of mice per group indicated.

**Figure 4.**
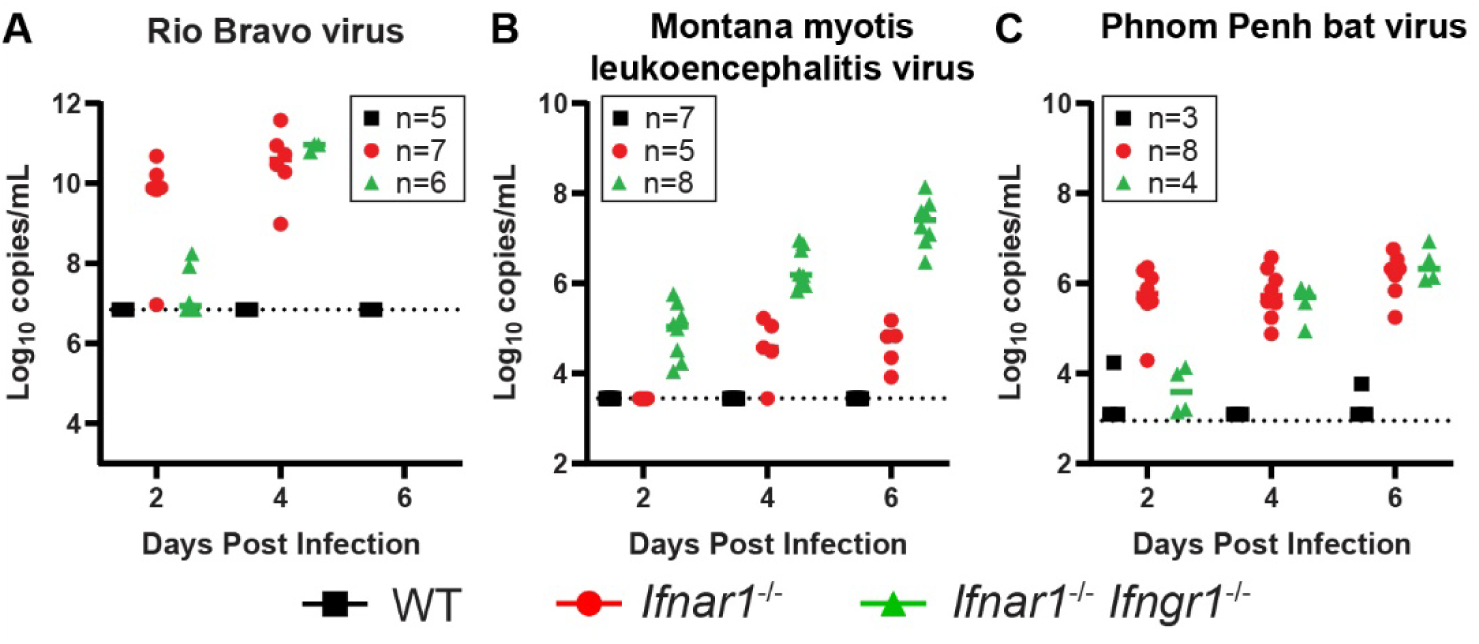
Detection of NKVFV viremia by RT-qPCR. Five to six-week-old male and female wild-type (WT), *Ifnar1*^−/−^, and *Ifnar1*^−/−^ *Ifngr1*^−/−^ mice were infected with 1000 FFU of the indicated flavivirus by subcutaneous inoculation in the footpad. **A.** Rio Bravo virus (RBV) (M64). **B.** Montana myotis leukoencephalitis virus (MMLV) (B 334 A-924 BF). **C.** Phnom Penh bat virus (PPBV) (A38-69). Mice were bled at 2-, 4-, and 6-days post-infection and viremia was measured by RT-qPCR. Data are combined from 2 to 3 experiments per virus, with the number of mice per group indicated.

### Bat-borne NKVFVs have variable replication kinetics in primary mouse cells

MMLV, PPBV, and RBV are bat-borne NKVFVs that cluster together phylogenetically (**Figure 1**). However, MMLV and PPBV did not cause lethality in *Ifnar1^−/−^* mice, compared to 100% lethality caused by RBV (**Figure 2**), with RBV also producing higher viremia **(Figure 3 and Figure 4)**. To evaluate replication of these three viruses in mouse cells, we generated mouse embryonic fibroblasts (MEF) from wild-type, *Ifnar1^−/−^*, and *Ifnar1^−/−^ Ifngr1^−/−^* mice and performed multi-step growth curves. MEFs were infected with RBV, MMLV, or PPBV at an MOI of 0.01. Supernatant was collected at 4, 24, 48, and 72 hours post-infection (hpi). Vero cells were included as a positive control for infection. We found that RBV was able to replicate in all cell types tested, with greater replication in *Ifnar1^−/−^*, and *Ifnar1^−/−^ Ifngr1^−/−^*MEFs compared to Vero cells and lower replication in wild-type MEF (**Figure 5A**). MMLV and PPBV exhibited greatly diminished replication compared to RBV, with MMLV replicating in *Ifnar1^−/−^* and *Ifnar1^−/−^ Ifngr1^−/−^* MEFs but not wild-type MEFs (**Figure 5B**), and almost no PPBV detected in any of the MEF lines (**Figure 5C**).

**Figure 5.**
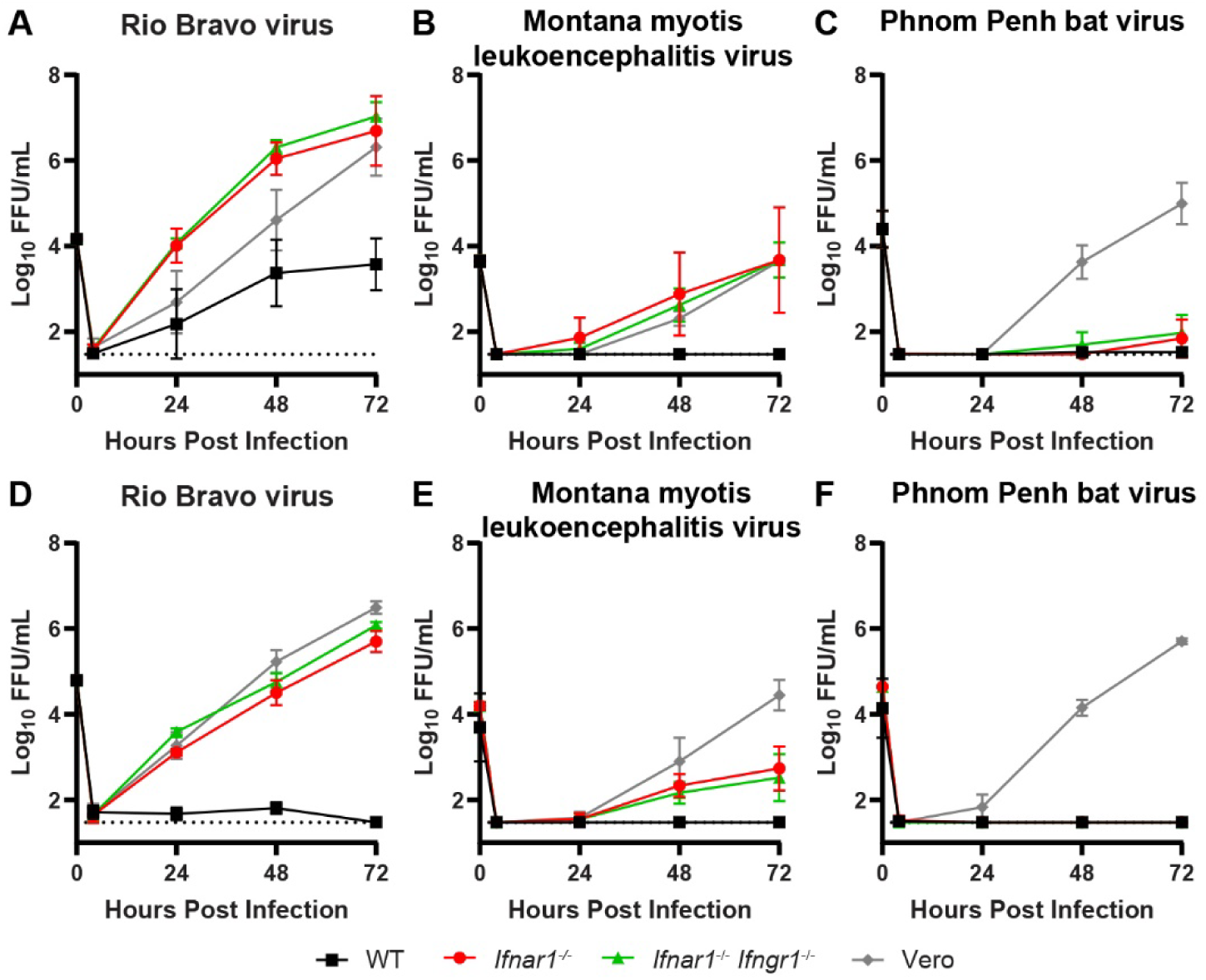
Bat-borne NKVFV have variable replication kinetics in mouse cells. Mouse embryonic fibroblasts (MEF) **(A-C)** and bone marrow-derived macrophages (BMDM) **(D-F)** were prepared from wild-type (WT), *Ifnar1*^−/−^, and *Ifnar1*^−/−^ *Ifngr1*^−/−^ mice. MEF, BMDM, and Vero cells were infected at an MOI of 0.01 with RBV, MMLV, or PPBV and infectious virus in the supernatant was quantified by FFA at 4-, 24-, 48-, and 72-hours post-infection. Data are shown as the mean ± SEM of 2 to 3 independent experiments performed in duplicate or triplicate.

Since flaviviruses typically target myeloid cells, rather than fibroblasts (40), we assessed NKVFV replication in bone marrow derived macrophages (BMDM) prepared from wild-type, *Ifnar1^−/−^*, and *Ifnar1^−/−^ Ifngr1^−/−^* mice. BMDMs and Vero cells were infected with RBV, MMLV, or PPBV at an MOI of 0.01 and supernatant was collected at 4, 24, 48, and 72 hpi. The overall trends in viral replication were similar in BMDMs compared to MEFs, with RBV replicating to high titers in *Ifnar1^−/−^* and *Ifnar1^−/−^Ifngr1^−/−^* BMDMs, while MMLV replicated only to low titers, and no infectious virus was detected in BMDM infected with PPBV (**Figure 5D-F**). In contrast to MEFs, where RBV and MMLV replicated to higher titers in *Ifnar1^−/−^Ifngr1^−/−^* cells compared to *Ifnar1^−/−^*, there was no significant difference in RBV or MMLV replication in *Ifnar1^−/−^Ifngr1^−/−^* BMDM compared to *Ifnar1^−/−^*. Overall, these results show that IFN-αβ signaling plays a key role in restricting RBV and MMLV replication in mouse cells. Strikingly, no PPBV replication was detected in *Ifnar1^−/−^Ifngr1^−/−^*MEFs or BMDMs, even though PPBV caused 100% lethality in *Ifnar1^−/−^Ifngr1^−/−^* mice, suggesting that other cell types may be key targets for PPBV infection.

### Bat-borne NKVFVs have variable replication kinetics in bat tissue-derived cell lines

RBV, MMLV, and PPBV have been isolated from wild-caught bats: RBV from salivary glands of *Tadarida brasiliensis*, MMLV from salivary glands of *Myotis lucifugus*, and PPBV from salivary glands and brown fat of *Cynopterus brachyotis* (**Table 2**) (35–37). To assess the ability of RBV, MMLV, and PPBV to replicate in bat-derived cell lines, we infected Tb1Lu (lung epithelial cells from *Tadarida brasiliensis*), Efk3B (kidney epithelial cells from *Eptesicus fuscus*, big brown bat), and LaBo (*Lasiurus borealis*, eastern red bat) cell lines with RBV, MMLV, or PPBV at an MOI of 0.01 and collected supernatants at 4, 24, 48, and 72 hpi. We found that RBV replicated to high titers in LaBo cells, equivalent to replication in Vero cells, and to lower titers in Efk3B and Tb1Lu cells (**Figure 6A**). MMLV replication in Efk3B and Tb1Lu cells was similar to MMLV replication in BMDM and MEF, being slow and first detected 48 hpi and increasing at 72 hpi. However, in Efk3B cells, MMLV replication peaked at 48 hpi then began to decline (**Figure 6B**). We did not detect PPBV replication in Tb1Lu or LaBo cells, concordant with a lack of PPBV replication in BMDM and MEF. However, PPBV did exhibit low level replication in Efk3B cells, which is the first example of PPBV replication in an immunocompetent cell line (**Figure 6C**). These results indicate that bat-borne NKVFV are able to infect bat-derived cell lines but the level of replication varies among viruses and cell lines.

**Figure 6.**
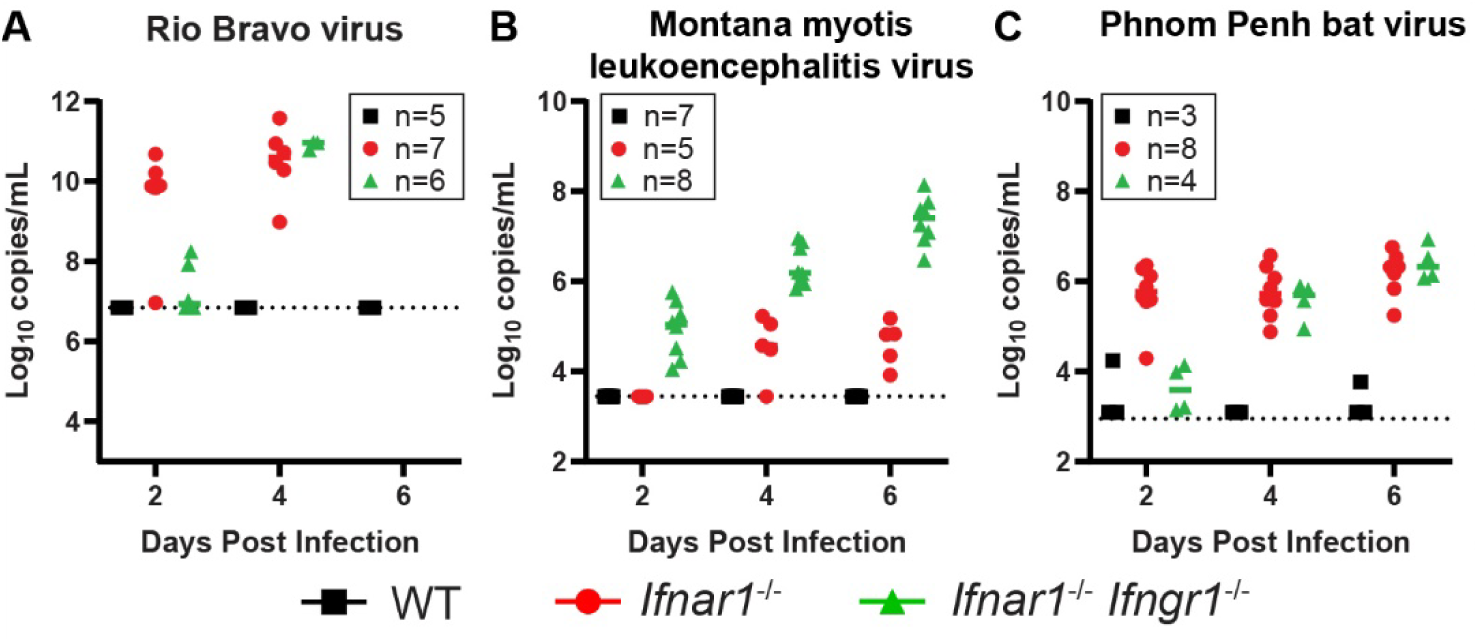
Bat-borne NKVFV have variable replication kinetics in bat-derived cell lines. TB1Lu (*Tadarida brasiliensis*), Efk3B (*Eptesicus fuscus*), LaBo (*Lasiurus borealis*), and Vero cells were infected with RBV, MMLV, or PPBV at an MOI of 0.01 and infectious virus in the supernatant was quantified by FFA at 4, 24, 48, and 72 HPI. Data are shown as the mean ± SEM of 2 to 3 independent experiments performed in duplicate or triplicate.

## DISCUSSION

Orthoflaviviruses can be categorized into four groups based on the role of arthropod vectors in their transmission: mosquito-borne, tick-borne, insect-specific, or no-known-vector flaviviruses; genetic relatedness generally corresponds to shared transmission mechanisms, but these groupings are not monophyletic (4–6). NKVFVs have been isolated from bats and rodents worldwide. Bat-borne NKVFVs have been isolated in Mexico, Trinidad, Uganda, Madagascar, Kyrgyzstan, Cambodia, Malaysia, Japan, and the United States from a range of species including both insectivorous bats (*Myotis lucifugus* (35)*, Molossus rufus* (37)*, Tadarida brasiliensis* (41)*, Chaerephon pumilus* (22, 23)*, Pteronotus parnellii* (37)*, Tadarida condylurus* (23), and *Miniopterus fuliginosus* (27)) and frugivorous bats (C*ynopterus brachyotis* (29, 36, 42)*, Eonycteris spelaea* (36), and *Macroglossus lagochilus* (29, 42)(2)). Rodent-borne NKVFVs have been isolated in the United States, Canada, Guatemala, and Japan from *Sigmodon hispidus* (29, 32, 43)*, Peromyscus maniculatas* (44, 45)*, Peromyscus leucopus* (32)*, Apodemus speciosus* (29), and *Clethrionomys rufocanus* (2, 29). More recently, additional orthoflaviviruses (and related unclassified members of the *Flaviviridae* family) have been identified from aquatic vertebrates and invertebrates by metagenomic methods or classical isolation from a sick animal in the case of salmon flavivirus (6, 46–50), altogether highlighting the wide range of hosts that harbor flaviviruses, beyond the well-studied vector-borne human pathogens.

Many important human pathogens belong to the mosquito-borne or tick-borne flaviviruses, with humans serving as amplifying hosts for some viruses and dead-end hosts for others (1, 7). Three NKVFVs have been reported to cause human disease: Rio Bravo virus, Apoi virus, and Modoc virus (2). RBV has caused one natural and seven laboratory-acquired infections, with presentation ranging from fever to central nervous system involvement (51). One laboratory-acquired APOIV infection has been reported, resulting in febrile illness, encephalitis, and bilateral leg paralysis (2). MODV was linked to aseptic meningitis in a child who handled a sick rodent (45).

In this study, we focused on three bat-borne NKVFVs: MMLV, RBV, and PPBV. These viruses are geographically distinct, with RBV and MMLV isolated from bats in the United States, and PPBV from Cambodia (2). Additionally, despite the importance of bats as reservoir hosts for emerging viruses (52–55), bat-borne flaviviruses have received relatively little attention compared to bat coronaviruses, paramyxoviruses, and filoviruses. Only a few NKVFVs have been investigated in pathogenesis studies and each study used a different animal model: Syrian hamsters (MODV) (9), SCID mice (MODV, MMLV) (13, 56), immunocompetent mice (MODV, RBV, MMLV, ENTV), outbred deer mice (MODV) (44), rhesus macaques (RBV) (51), and Leschenault’s rousette bats (YOKV) (10). Here we investigated NKVFV pathogenesis in wild-type, *Ifnar1*^−/−^, and *Ifnar1*^−/−^ *Ifngr1*^−/−^ mice, using a subcutaneous inoculation route, facilitating comparisons between NKVFVs and other flaviviruses which are commonly studied in these models. We chose a subcutaneous inoculation route because this route commonly is used for studies of vector-borne flaviviruses, but the transmission routes of NKVFVs remain unknown. Future studies could investigate how different inoculation routes impact susceptibility to NKVFV infections. Likewise, our viral load measurements assessed viremia, a key outcome of infection with vector-borne flaviviruses, but viremia may not be important for viruses that are not transmitted by hematophagous arthropod vectors; future studies could measure viral loads in epithelial tissues that might be more relevant to other transmission routes. The pathogenic phenotypes of NKVFVs in these mouse models lay the foundation for future studies investigating NKVFV-specific and pan-flavivirus pathogenic mechanisms and evaluating the breadth of action of candidate vaccines and therapies against flaviviruses, as well as providing experimental models at the ready to respond to future emerging flaviviruses.

## MATERIALS AND METHODS

### Mice

All animal husbandry and experimental protocols were approved by the University of North Carolina at Chapel Hill Institutional Animal Care and Use Committee. Five- to six-week-old male and female wild-type, *Ifnar1^−/−^,* or *Ifnar1^−/−^ Ifngr1^−/−^* mice on a C57BL/6 background were used. Mice were inoculated with 1,000 FFU of virus in a volume of 50µL subcutaneously in the footpad and monitored daily for disease signs for 21 days. Mice were euthanized upon reaching humane endpoints, including nonresponsiveness, or severe neurologic disease signs (hunching, paralysis) that interfered with the ability to access food and water. To assess viremia, blood was collected 2, 4, and 6 dpi via submandibular bleed with a 5mm Goldenrod lancet into serum separator tubes (BD). Serum was separated by centrifugation for 5 minutes at 15,000 rpm and stored at −80° for quantification of virus by focus-forming assay or RT-qPCR.

### Cells

Vero (African green monkey kidney epithelial) cells were maintained in Dulbecco’s modified Eagle medium (DMEM) containing 5% heat-inactivated fetal bovine serum (FBS), HEPES buffer, and L-glutamine at 37°C with 5% CO2. Immortalized *Eptesicus fuscus* kidney epithelial (Efk3B) cells (Kerfast, catalog #ESA001) were maintained in DMEM containing 10% heat-inactivated FBS and 1x HEPES buffer. *Lasiurus borealis* (LaBo) cells were maintained in DMEM containing 10% heat-inactivated FBS. Efk3B and LaBo cell lines were provided by Dr. Ralph Baric (UNC). *Tadarida brasiliensis* lung epithelial (Tb1Lu) cells were maintained in DMEM containing 10% heat-inactivated FBS.

### Primary cells

Mouse embryonic fibroblasts (MEFs) were prepared from E15 embryos. Embryos were removed from the gravid uterus, placed in PBS, decapitated, and gut and liver were removed. Embryos were then minced with scalpels, trypsinized (1 ml per embryo), pipetted up and down with a 10 ml serological pipette to break up any chunks, and incubated for 5-10 min at room temperature. Cells were resuspended in DMEM supplemented with non-essential amino acids, L-Glutamine, Pen/Strep, and 10% heat-inactivated FBS and then pelleted by centrifugation at 1000 rpm for 5 min at 4°C. Supernatants were removed and cell pellets were resuspended in fresh media and pelleted again by centrifugation at 1000 rpm for 5 min at 4°C. Cell pellets were resuspended in 1 ml per embryo of fresh media and plated into culture flasks (1.5 embryos per T-150 flask) in 25 ml of fresh media and incubated at 37°C with 5% CO_2_. After 24 hours, media was removed, cells were washed with 1X PBS, and fresh media was added. When monolayers reached near-confluency, MEFs were frozen down in DMEM supplemented with non-essential amino acids, L-Glutamine, Pen/Strep, 30% heat-inactivated FBS, and 20% DMSO and stored in liquid nitrogen. Thawed MEFs were seeded in 6-well plates at 2 × 10^5^ cells per well in DMEM supplemented with non-essential amino acids, L-Glutamine, Pen/Strep, and 10% heat-inactivated FBS. Bone marrow-derived macrophages (BMDM) were generated from wild-type, *Ifnar1^−/−^,* or *Ifnar1^−/−^Ifngr1^−/−^* mice on a C57BL/6 background. Mice were euthanized and femurs and tibias were isolated from hind limbs. Bone marrow was flushed out with 10 ml DMEM delivered via syringe with 25G ½ inch needle. Bone marrow was pooled and pipetted up and down with a 5 ml serological pipette to break up large chunks. Cells were pelleted by centrifugation at 1500 rpm for 5 min at 4°C. Supernatants were removed and cell pellets were resuspended in ACK Red Blood Cell Lysis Buffer containing 150 mM NH_4_Cl, 10 mM KHCO_3_, and 0.1 mM EDTA pH 7.3, and incubated for 2-3 minutes. Cells were resuspended in DMEM containing 10% heat-inactivated FBS and then pelleted by centrifugation at 1500 rpm for 5 min at 4°C. Cell pellets were resuspended in DMEM containing 10% heat-inactivated FBS and counted. 12-well non-TC treated plates were seeded with 1.5 × 10^5^ cells/well in 1 ml of DMEM containing L-Glutamine, NaPyr, Pen/Strep, 10% heat-inactivated FBS, and 40 ng/ml mouse M-CSF (BioLegend 576406) and incubated for 7 days at 37°C with 5% CO_2_.

### Viruses

All experiments in this study were conducted under BSL-2 containment, under protocols approved by the UNC Institutional Biosafety Committee and Environmental Health and Safety. Most virus isolates were obtained from the World Reference Center for Emerging Viruses and Arboviruses: Culex flavivirus (CXFV, HOU24518)(57), Aedes flavivirus (AEFV, strain MO-M6)(3), Aroa virus (AROAV, strain Maracay 01809)(58), Banzi virus (BANV, strain SA H 336)(59), Iguape virus (IGUV, strain SP AN 71686)(60), Entebbe bat virus (ENTV, strain IL 30)(22), Sokuluk virus (SOKV, strain LEIV 400K)(26), Yokose virus (YOKV, strain Oita 36)(27, 61), Apoi virus (APOIV, strain Apoi)(31), Modoc virus (MODV, strain M 544)(30), Jutiapa virus (JUTV, strain JG 128)(61), Sal Vieja virus (SVV, strain 78 TWM 106)(61), Kadam virus (KADV, strain T100)(62, 63), Langat virus (LGTV, strain TP-21)(64), Dakar bat virus (DBV, strain BP180 (33), strain AD D249 (65), and strain IBAN 8646 (66)), Usutu virus (USUV, strain SA AR 1776)(67), Kedougou virus (KEDV, strain DAK AR D 14701)(61), Montana myotis leukoencephalitis virus (MMLV, strain B 334 A-924 BF)(35), Rio Bravo virus (RBV, strain M64)(37), Phnom Penh bat virus (PPBV, strain A38-69)(36), and Tamana bat virus (TABV, strain TR 127154)(37). Zika virus (ZIKV, strain H/PF/2013)(68) was obtained from the US Centers for Disease Control and Prevention. Virus stocks were passaged twice in Vero cells in DMEM supplemented with 5% heat-inactivated FBS, 1x HEPES (10mM based on 5 mL of 1 M HEPES into 500 mL DMEM), and 1x Glutamax (10mM based on 5 mL of 1 M Glutamax into 500 mL DMEM) and titered by focus forming assay (FFA).

### Viral focus-forming assay

Virus quantification was performed by FFA on Vero cells. Duplicates of serial 10-fold dilutions of virus in viral growth medium (DMEM containing 2% FBS and 20 mM HEPES) were applied to Vero cells in 96-well plates and incubated as described above for 1 h. Following virus adsorption, the monolayers were overlaid with 1% methylcellulose in Eagle minimum essential medium (MEM). Plates were incubated at 37°C with 5% CO_2_ for virus-specific incubation times (**Table 3**). Following fixation with 2% paraformaldehyde for 1 h at room temperature, plates were incubated with 100 ng/ml of flavivirus-cross-reactive human monoclonal antibody (MAb) PB-006-01 for 2 h at room temperature or overnight at 4°C. After incubation at room temperature for 2 h with a 1:5,000 dilution of horseradish peroxidase (HRP)-conjugated goat anti-mouse IgG (Sigma), foci were detected by the addition of TrueBlue substrate. Foci were quantified with an Immunospot analyzer (Cellular Technology Limited).

**Table 3:**
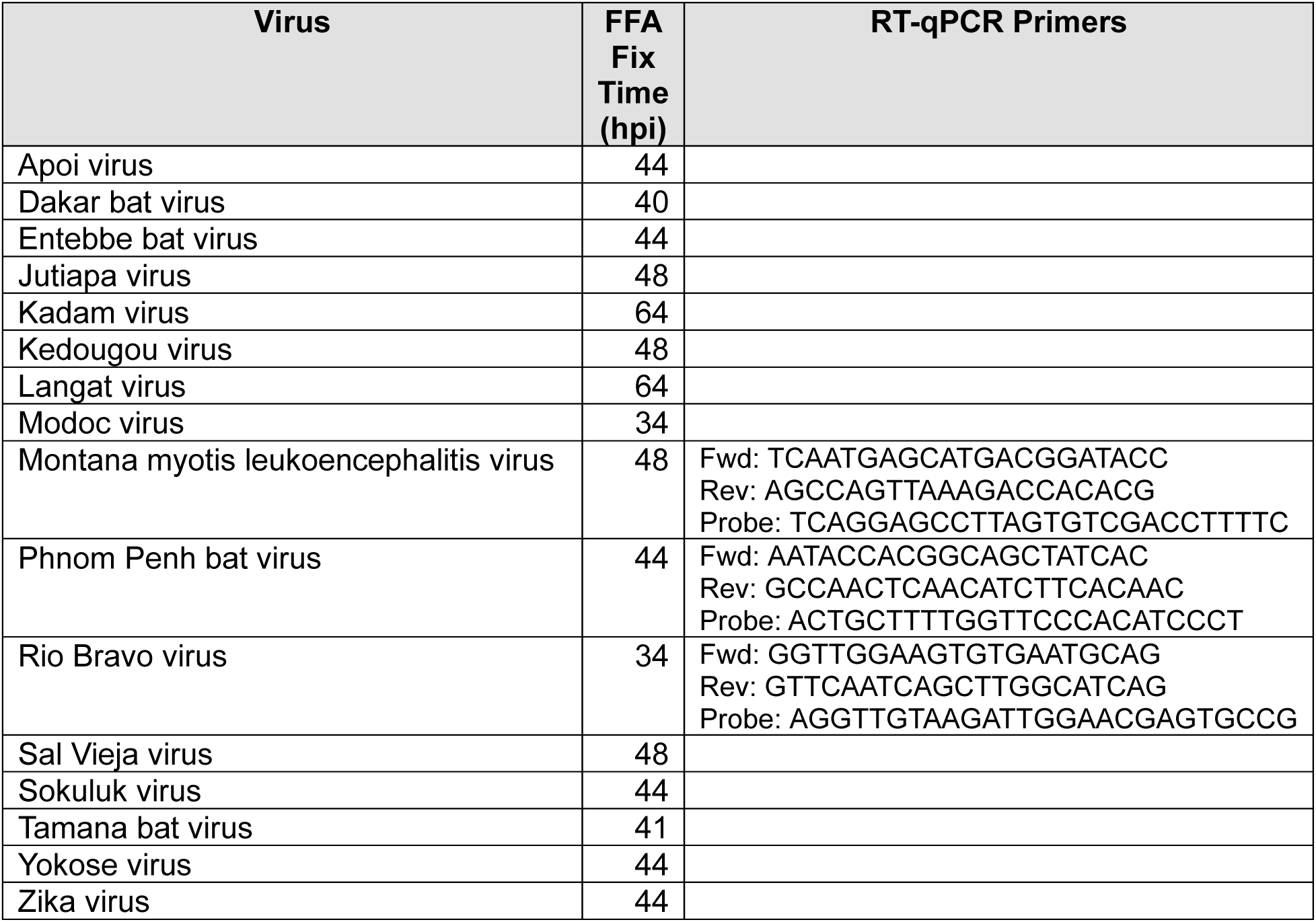
Focus-forming assay conditions and RT-qPCR primers.

### Measurement of viremia by RT-qPCR

RNA was extracted from serum using the Viral RNA Mini Kit (Qiagen). Viral RNA levels were determined via TaqMan one-step RT-qPCR using the iTaq - Universal Probes One-Step Kit (BioRad) on a CFX96 Touch Real-Time PCR Detection System (Bio-Rad), using standard cycling conditions. Viremia is expressed on a Log_10_ scale as copies per mL of serum, based on standard curves produced using serial 10-fold dilutions of plasmids containing a 779 bp sequence from NS5 of MMLV, RBV, or PPBV (Twist Bioscience). The indicated primers and probes (**Table 3**) were purchased from Integrated DNA Technologies and were used at 500 nM.

### Viral growth curves

Vero, BMDM, MEF, Efk3B, LaBo, and Tb1Lu cells were infected at a multiplicity of infection (MOI) of 0.01 and incubated at 37°C with 5% CO_2_ for 1 hour. After, the media was removed, the wells were washed with PBS, and fresh media was added. Supernatant was collected at 4, 24, 48, and 72 hours post-infection and stored at −80°C until analysis by FFA on Vero cells.

### Deep sequencing of viral stocks

#### RNA extraction and cDNA synthesis

1 mL of clarified viral supernatant was divided into 3 equal aliquots. RNA was extracted from each aliquot with TRIzol LS and co-precipitated with glycogen (ThermoFisher) according to the manufacturer’s protocol. Each set of 3 pellets was re-pooled and dissolved in RNase-free water. First strand cDNA was generated from each RNA sample using SuperScript IV reverse transcriptase (ThermoFisher). Second strand synthesis was performed with the Second Strand Synthesis module (NEB) and purified with a Monarch DNA Cleanup Kit (NEB).

#### Library preparation and sequencing

1 ng of purified double stranded viral cDNA was used to Tagment, amplify, and cleanup sequencing libraries with the Nextera XT kit (Illumina) according to the manufacturer’s protocol. Library normalization and pooling was performed by the UNC High Throughput Sequencing Facility. Normalized libraries were sequenced on a HiSeq platform (Illumina) with paired end 150 bp read length at the UNC High Throughput Sequencing Facility.

#### Sequence processing and analysis

Viral-enriched RNA-seq reads were first assembled de novo using Trinity to generate an initial reference sequence. The sequence was then refined through iterative consensus calling using a multi-strategy approach: viral reads were aligned back to the reference using both conservative BWA-MEM (v0.7.17) parameters and sensitive minimap2 (v2.26) alignments, followed by variant calling with bcftools mpileup/call (v1.22) and consensus generation with bcftools consensus. Uncertain 3′ regions were further resolved using SPAdes (v4.2.0) local assembly and BLAST (v2.16.0) analysis to identify potential extensions, with seqtk (v1.5) used for sequence manipulation and samtools (v1.23) for BAM file processing throughout. For other flavivirus samples with established reference genomes available, a simplified workflow was followed using direct alignment of viral-enriched reads to known NCBI reference strains using BWA-MEM (v0.7.17) or minimap2 (v2.26), followed by standard samtools/bcftools consensus calling to generate sample-specific consensus sequences. A read depth of greater than 50x was required for consensus calling in all situations.

#### Phylogenetic analysis

Full polyprotein amino acid sequences were downloaded from Genbank or from our own deep sequencing results. Sequences were aligned using MUSCLE through the Geneious interface. Alignments were exported as FASTA files and a maximum likelihood phylogenetic tree was estimated using IQtree at the command line. The phylogenetic tree was visualized using FigTree v1.4.4. (http://tree.bio.ed.ac.uk/software/figtree/). Sequences were deposited in GenBank (**Table 1**). Silhouette images are from PhyloPic.org.

## ACKNOWLEDGEMENTS

This work was supported by R01 AI170625 (H.M.L.) and U19AI171292 (W.A.S. and N.J.M.). H.M.L. holds an Investigators in the Pathogenesis of Infectious Disease Award from the Burroughs Wellcome Fund. M.E.D. and J.R.N. were supported by T32 AI007419.

